# Persistent DNA Hyper-methylation Dampens Host Anti-Mycobacterial Immunity

**DOI:** 10.1101/805457

**Authors:** Andrew R. DiNardo, Kimal Rajapakshe, Tomoki Nishiguchi, Godwin Mtetwa, Qiniso Dlamini, Jaqueline Kahari, Sanjana Mahapatra, Alexander Kay, Gugu Maphalala, Emily Mace, Sandra L. Grimm, George Makedonas, Jeffrey D Cirillo, Mihai G. Netea, Reinout van Crevel, Cristian Coarfa, Anna M. Mandalakas

**Affiliations:** The Global Tuberculosis Program, Texas Children’s Hospital, Immigrant and Global Health, Department of Pediatrics, Baylor College of Medicine, Houston, USA; Dan L Duncan Comprehensive Cancer Center, Baylor College of Medicine, Houston, USA; Baylor-Swaziland Children’s Foundation, Mbabane, Swaziland; Ministry of Health, Mbabane, Swaziland; Department of Pediatrics, Columbia University Medical Center, New York, NY, USA; JES Tech, Department of Immunology, NASA, Houston, TX, USA; Department of Microbial Pathogenesis and Immunology, Texas A&M Health Science Center, Bryan, TX, USA; Department of Internal Medicine and Radboud Center for Infectious Diseases, Radboud University Medical Center, Nijmegen, Netherlands

**Keywords:** Tuberculosis, DNA methylation, immune exhaustion

## Abstract

*Mycobacterium tuberculosis (Mtb*) has co-evolved with humans for millennia and developed multiple mechanisms to evade host immunity. Restoring host immunity in order to shorten existing therapy and improve outcomes will require identifying the full complement by which host immunity is inhibited. Perturbing host DNA methylation is a mechanism induced by chronic infections such as HIV, HPV, LCMV and schistosomiasis to evade host immunity. Here, we evaluated the DNA methylation status of TB patients and their asymptomatic household contacts demonstrating that TB patients have DNA hyper-methylation of the IL-2-STAT5, TNF-NF-ϰB and IFN-γ signaling pathways. By MSRE-qPCR, multiple genes of the IL-12-IFN-γ signaling pathway (*IL12B, IL12RB2, TYK2, IFNGR1, JAK1* and *JAK2*) were hyper-methylated in TB patients. The DNA hyper-methylation of these pathways is associated with decreased immune responsiveness with decreased mitogen induced upregulation of IFN-γ, TNF, IL-6 and IL-1β production. The DNA hyper-methylation of the IL-12-IFN-γ pathway was associated with decreased IFN-γ induced gene expression and decreased IL-12 inducible up-regulation of IFN-γ. This work demonstrates that immune cells from TB patients are characterized by DNA hyper-methylation of genes critical to mycobacterial immunity resulting in decreased mycobacteria-specific and non-specific immune responsiveness.

## Introduction

Tuberculosis (TB) continues to be a global problem with 10 million cases and over a million deaths each year^1^. Anti-TB therapy requires six months, despite the majority of individuals becoming culture negative within the first two months of therapy. It is unique to mycobacterial disease for therapy to be required for so many months after culture conversion. While multiple mechanisms of TB-induced immunosuppression have been identified, it is unknown why anti-TB therapy must be continued for months after an individual becomes culture negative. Further, after successful TB therapy, patients have a 13-fold increased risk of disease recurrence compared to the general population^2^. Recurrent disease could be related to strain virulence, environmental risk factors or persistent perturbations of host immunity. It is therefore critical to identify mechanisms of host immunosuppression that persist despite successful TB therapy.

TB has been recognized as an archetypical chronic infection since at least 1882. Since 1984, the Clone 13 strain of murine LCMV has emerged as the prototypical animal model of chronic infection^3^. Chronic infection with clone 13 LCMV induces immune exhaustion, which is defined by decreased antigen induced immune cell proliferation, decreased immune effector function and increased expression of immune checkpoint inhibitors^4-6^. Immune exhaustion is mediated by epigenetic alterations, conformational changes in immune cell DNA methylation and chromatin architecture that inhibits robust cell-mediated immunity^7^. These epigenetic changes are driven by DNA methyltransferase (DNMT) and the Enhancer of Zeste 2 Polycomb Repressive complex (EZH2) that imbues a repressive chromatin state, resulting in an immune non-responsive state^8-11^. DNA methylation is a common and stable epigenetic alteration that blocks transcription factor binding and inhibits gene expression and mediates immune exhaustion. Once exhausted, the epigenetic perturbations persist and the immune phenotype remains subdued even if cells are rested for three weeks, or if they are transferred into healthy, non-infected mice^10,12^. Similarly, humans chronically infected with HIV or schistosomiasis have DNA methylation perturbations of immunity that persist for months to years after successful therapy^8,13,14^.

Given that chronic antigen stimulation from HIV, schistosomiasis, or LCMV induces long-lasting epigenetic-mediated immune exhaustion, we were concerned that TB disease may also result in persistent detrimental epigenetic changes^8,10,13-15^. We hypothesized that TB patients would have a DNA methylation landscape similar to chronic LCMV. Mouse models of chronic LCMV have demonstrated persistent epigenetic perturbations, with hypomethylating agents capable of restoring host immunity^9,10,12^. We thought it pertinent to investigate the longitudinal DNA methylation of host immune cells before and after successful anti-TB therapy. In a cohort of microbiologically-confirmed adults with pulmonary TB from Eswatini, TB patients exhibit a hypo-responsive immune phenotype with decreased immune proliferation and decreased mycobacterial and mitogen induced cytokine production in association with DNA hyper-methylation of multiple critical immune genes and pathways. These DNA methylation perturbations persisted 6 months after successful anti-TB therapy and identifies a plausible mechanisms of suppressed host immunity.

## Results

We previously described a cohort of asymptomatic, TB-exposed children from Eswatini (formerly known as Swaziland), with schistosomiasis induced DNA hyper-methylation and associated inhibition of BCG and *Mycobacterium tuberculosis* (*Mtb)* immunity^14^. Here, to evaluate if TB induces similar DNA methylation perturbations that correspond with immune exhaustion, we assessed adults with TB symptoms and microbiologically-confirmed (by culture and/ or Gene Xpert) pulmonary TB. We compared their immune phenotype and DNA methylation status to their asymptomatic, healthy household contacts who remained asymptomatic for 12 months after initial exposure (Fig.1). All study participants were BCG vaccinated, as determined either via vaccine records and/ or BCG scar. HIV screen was performed at time of study initiation and then annually. TB treatment outcomes were defined by both WHO criteria and a simplified clinically relevant definition^16^.

**Figure 1:**
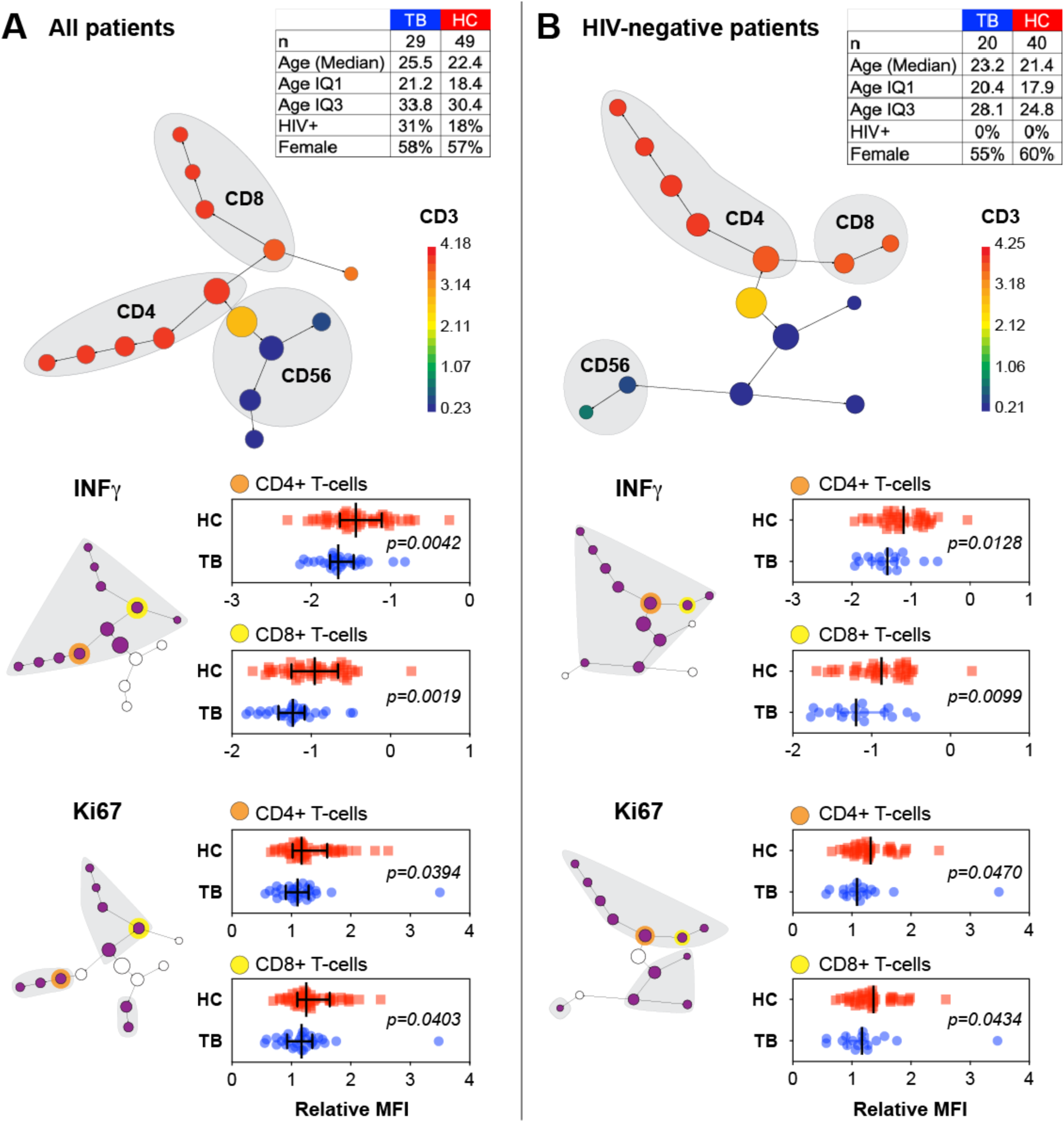
TB patients exhibit an exhausted immune phenotype. PBMCs from individuals with Tuberculosis (TB; n = 29) and their asymptomatic household contacts (HC; n = 49) underwent antigenic stimulation with BCG sonicate followed by flow-cytometry based multi-dimensional immune profiling. CITRUS (cluster identification, characterization, and regression) identified CD4, CD8, CD3 and CD56 subsets. We further defined NK cells as CD3^-^CD56^+^ cells. (**A**) Citrus clustering identifies CD4, CD8 and NK cell populations with CD3 signal depicted by heatmap. Grey-shaded sub-clusters depict decreased IFN-γ and Ki-67 among Helper T cells (CD3^+^CD4^+^), CTLs (CD3+CD8+) and NK (CD3^-^CD56^+^) cells. Representative bar graphs of the MFI clusters are depicted from the yellow highlighted cluster. (**B**) Re-analysis of CITRUS with exclusion of HIV co-infected individuals of TB (n = 20) and controls (n = 40) depicts CD4, CD8 and CD56 lymphocyte subsets with decreased Ki-67 and IFN-γ (grey-shaded clusters have decreased IFN-γ and Ki-67; representative bar graphs of MFI clusters depicted).

### TB patients demonstrate immune exhausted phenotypes

To evaluate the cell-specific and mycobacterial-specific immunity of TB patients, peripheral blood mononuclear cells (PBMCs) were stimulated with ESAT-6 and CFP-10 (*Mtb*-specific antigens) or BCG sonicate followed by flow cytometry based multi-dimensional immune profiling (MDIP). Cells were stained subsequently for viability, cell surface and intracellular markers. Multi-dimensional immune changes were evaluated using CITRUS (clustering identification, characterization and regression^17^), identifying subsets of helper T cells (CD3^+^CD4^+^), cytotoxic lymphocytes (CD3^+^CD8^+^; CTLs) and natural killer cells (CD3^-^CD56^+^; NK) (Fig 1). Median fluorescent intensity (MFI) of each cluster was compared between Healthy controls (HCs) and individuals with TB. CITRUS identified that compared to controls, participants with TB have Helper T cells, CTLs and NK cells with decreased IFN-γ and proliferative capacity in response to both *Mtb*-specific antigen (ESAT6 and CFP-10) and BCG (Fig 1a; Supplemental Fig. 1a). Compared to controls, individuals with TB were more likely to be co-infected with HIV, however after excluding people living with HIV (PLWH), participants with TB still have lymphocytes (CD4^+^, CD8^+^ and CD3^-^CD56^+^) with decreased IFN-γ and proliferative capacity (Fig. 1b; Supplemental Fig. 1). Study participants with TB, and not HIV co-infected, had an 21% and 25% decreased BCG induced IFN-γ production compared to HIV-uninfected controls, respectively for CD4 and CD8 populations. Similarly, HIV uninfected TB participants had an 16% and 14% decreased Ki-67, respectively for CD4 and CD8 populations.

The immune exhausted phenotype was not mycobacteria-specific as the decreased proliferative capacity also present after stimulation with the super-antigen staphylococcal enterotoxin B (SEB; Supplemental Fig. 1b). Increased expression of immune checkpoint blockade, as measured by PD1, CTLA4 and/or TIM3 is another characterization of immune exhaustion and similar to previous reports^18,19^, compared to asymptomatic, HCs, TB patients had an increased abundance of PD1 expressing NK cells (Supplemental Fig. 1c). A recent report demonstrated that TB-exposed individuals who did not progress to active disease had increased abundance of cytotoxic NK cells^20^. Only household contacts that remained asymptomatic and did not progress to active TB disease for 12 months were included in this analysis and similar to the recent report, household contacts in this cohort that did not progress had increased IFN-γ, T-bet and perforin production from NK cells compared to individuals with TB (Fig 1b; Supplemental Fig. 1). In contrast, TB patients had CTLs and helper T cells with increased perforin production (Supplemental Fig. 1).

### The TB DNA methylome resembles the immune exhaustion epigenetic landscape

Epigenetic mechanisms mediate immune exhaustion and therefore we evaluated the global DNA methylation status of TB participants using the EPIC array. From bulk PBMCs, cell-specific DNA methylation was determined using epigenetic deconvolution (EDEC)^21^. EDEC identified global differential methylation changes in Helper T cells, CTLs, NK cells and monocytes both at baseline and 6 months after completion of successful ATT, 12 months after study enrollment (Fig. 2d). The IL-2-STAT5 pathway, a critical component of cell proliferation, was differentially methylated at baseline and 6 months after successful therapy in all lymphocyte subsets. The PI3K-AKT pathway, which modulates the intracellular signaling of the immuno-metabolic pathway in both adaptive and innate cells, was differentially methylated in all cell types except CD4^+^ T cells. Similarly, the TNF-NFϰB signaling pathway was differentially methylated in all evaluated cell types except Helper T cells. IFN-γ signaling, critical to anti-mycobacterial immunity, was differentially methylated in all cell types at baseline and 6 months after the completion of ATT (Fig 2b). Specifically, the lymphocyte subsets (CD4^+^ Helper T cells, CD3^+^CD8^+^ CTLs and CD3^-^CD56^+^ NK cells) demonstrated DNA hyper-methylation the IFN-γ signaling pathway, while this pathway was hypo-methylated in monocytes (Fig 3).

**Figure 2:**
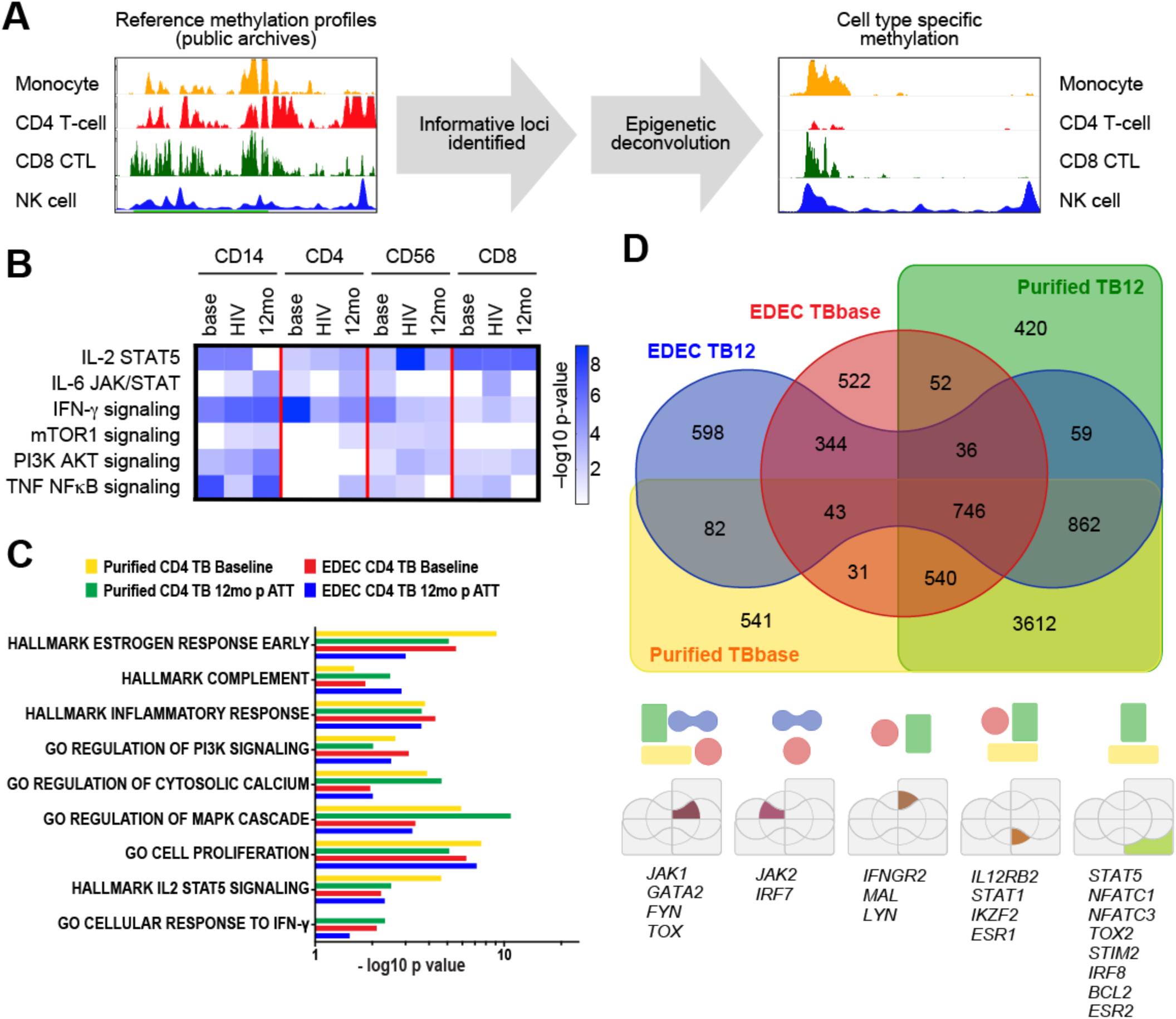
Global DNA methylation perturbations persist 6-months after successful anti-TB therapy. The Illumina DNA Methylation EPIC array was performed on bulk peripheral blood mononuclear cells (PBMCs) from asymptomatic household contacts (n = 10), individuals with TB (n = 15; TB/HIV^-^ = 7; TB/HIV^+^ = 8) at baseline. All individuals with TB had treatment success. For individuals without HIV co-infection, DNA methylation status was evaluated at baseline and 12-months later, 6 months after completion of successful anti-TB therapy (ATT). (**A**) To ascertain the immune cell-specific DNA methylation changes, cell-specific DNA methylation reference profiles were downloaded from public archives, informative loci were identified, and cell type specific methylation profiles were identified. (**B**) Cell-specific DNA methylation differences are shown from TB patients and TB/HIV patients at baseline and TB patients 6 months after completion of successful anti-TB therapy. All results are compared to healthy controls. GO pathway analysis for monocytes (CD14^+^), helper T cells (CD3^-^CD4^+^), NK cells (CD3^-^CD56^+^) and CTLs (CD3^+^CD8^+^). (**C-D**) DNA methyl EPIC results at the gene and pathway level from EDEC identified helper T cells (CD3^+^CD4^+;^ n = 8 with TB and 10 controls) and purified helper T cells (CD3^+^CD4^+^; n = 2) is compared.

**Figure 3:**
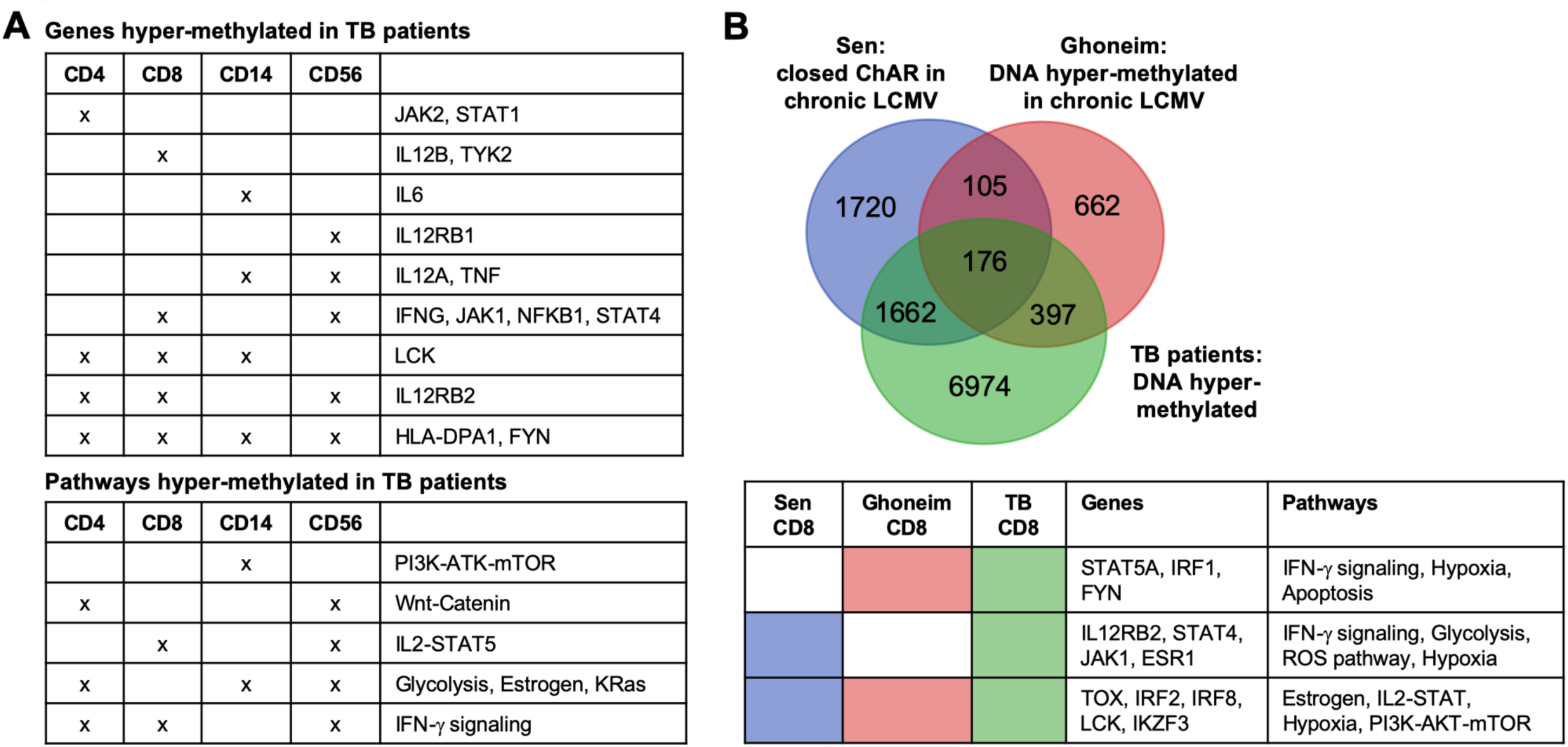
The TB epigenetic landscape is similar to LCMV-induced immune exhaustion. (**A**) DNA hyper-methylation of genes and GO pathways depicted by Venn diagrams of the CD14^+^ monocytes, CD4^+^ helper T cells, CD3^-^CD56^+^ NK cells and CD8^+^ cytotoxic T cells (CTLs). (**B**) Genes of CD8+ CTLs with DNA hyper-methylation by TB patients from this cohort were compared to genes with DNA hyper-methylation (Ghoneim et al) and closed chromatin accessible regions (ChAR; Sen et al) induced by chronic LCMV-induced immune exhaustion.

To validate the deconvolution results, CD4^+^ T cells were isolated and DNA methylation evaluated at baseline and 6 months after successful ATT. Of note, cell proliferation, intracellular calcium signaling, IL-2-STAT5 and the IFN-γ pathways were hyper-methylated in both isolated CD4^+^ T cells and the deconvoluted CD4^+^ T cells (Fig. 2c). Analysis of isolated CD4^+^ T cells confirmed results from EDEC for 82% of hypermethylated genes (probability of overlap =9.3 x10^−5^). Baseline DNA hypermethylation, as identified by both EDEC and in isolated CD4^+^ T cells was found in *JAK1, IL12RB2, STAT1, FYN, GATA2, IKZF2 and TOX* (Fig 2d).

Previous studies of chronic LCMV, cancer, HIV and schistosomiasis identified persistent epigenetic-mediated immune exhaustion long after removal of the chronic antigen stimulation^8,10,13,14^. The DNA hyper-methylation in TB patients, both at baseline and six months after successful anti-TB therapy, resembled the closed chromatin confirmation induced by chronic infection LCMV models. Similarities between the epigenetic landscape of murine LCMV and TB patients was evaluated by comparing the closed chromatin accessible regions (ChAR) previously published by Sen et al^4^ and DNA hyper-methylated regions previously published by Ghoneim et al^9^ from chronic LCMV models to the CD8 DNA hyper-methylated regions of TB patients from this study. In particular, DNA hyper-methylation and closed chromatin conformation of the IFN-γ pathway (*IL12RB2, JAK1, STAT4*) was seen both in TB and chronic LCMV (Fig.3b; probability of overlap <10^−5^). In addition to DNA methylation changes being induced by chronic antigenic exposure, *Mtb* itself releases epigenetic-modifying enzymes that target host chromatin and DNA methylation. For example, Rv2966c is a DNA methyltransferase that is secreted by *Mtb* and chaperoned into host immune cell nuclei^22^. In *in vitro* models, Rv2966c induces DNA hyper-methylation of host immune cells^23^ in a manner similar to that seen in this cohort of TB patients, including hyper-methylation of *IL12RB2, STAT4, IFNG, IRF1* and *JAK1* (Supplemental Figure 2).

### TB patients exhibit DNA hyper-methylation of the IL-12-IFN-γ signaling pathway

To validate the DNA methylome results, targeted MSRE-qPCR was implemented. While the EPIC array evaluates DNA methylation via bisulfite conversion, MSRE quantifies methylation status using ***m***ethylation ***s***pecific ***r***estriction ***e***ndonuclease isochizomers. MSRE-qPCR of isolated CD4^+^ T cells confirmed DNA hyper-methylation of the IL-12-IFN-γ pathway (Fig.4; n = 11). Specifically, *IL12B* demonstrated 6.1-fold increased methylation (p = 0.007); *IL12RB2* was hyper-methylated 35.2-fold (p < 0.03); *TYK2* demonstrated 17.7-fold increased methylation (p = 0.008); *IFNGR1* was hyper-methylated 24.0-fold (p = 0.007); *JAK1* and *JAK2* demonstrated 33.3 and 11.7-fold increased DNA methylation (p = 0.004 and 0.03), respectively (Fig. 4). *STAT1* and *IRF1* were not statistically hyper-methylated (p = 0.21 and 0.15, respectively; Fig. 4b). Regulation of immunity is epigenetically controlled at specific pathways, but also by a network of transcription factors^24^. By MSRE-qPCR, the CD4^+^ T cells from TB patients demonstrated DNA hyper-methylation of the transcription factors *NFATC1, BATF3, ID2, PPARG, RUNX2, IRF5 and IKZF1* (p< 0.001; Fig. 4c).

**Figure 4:**
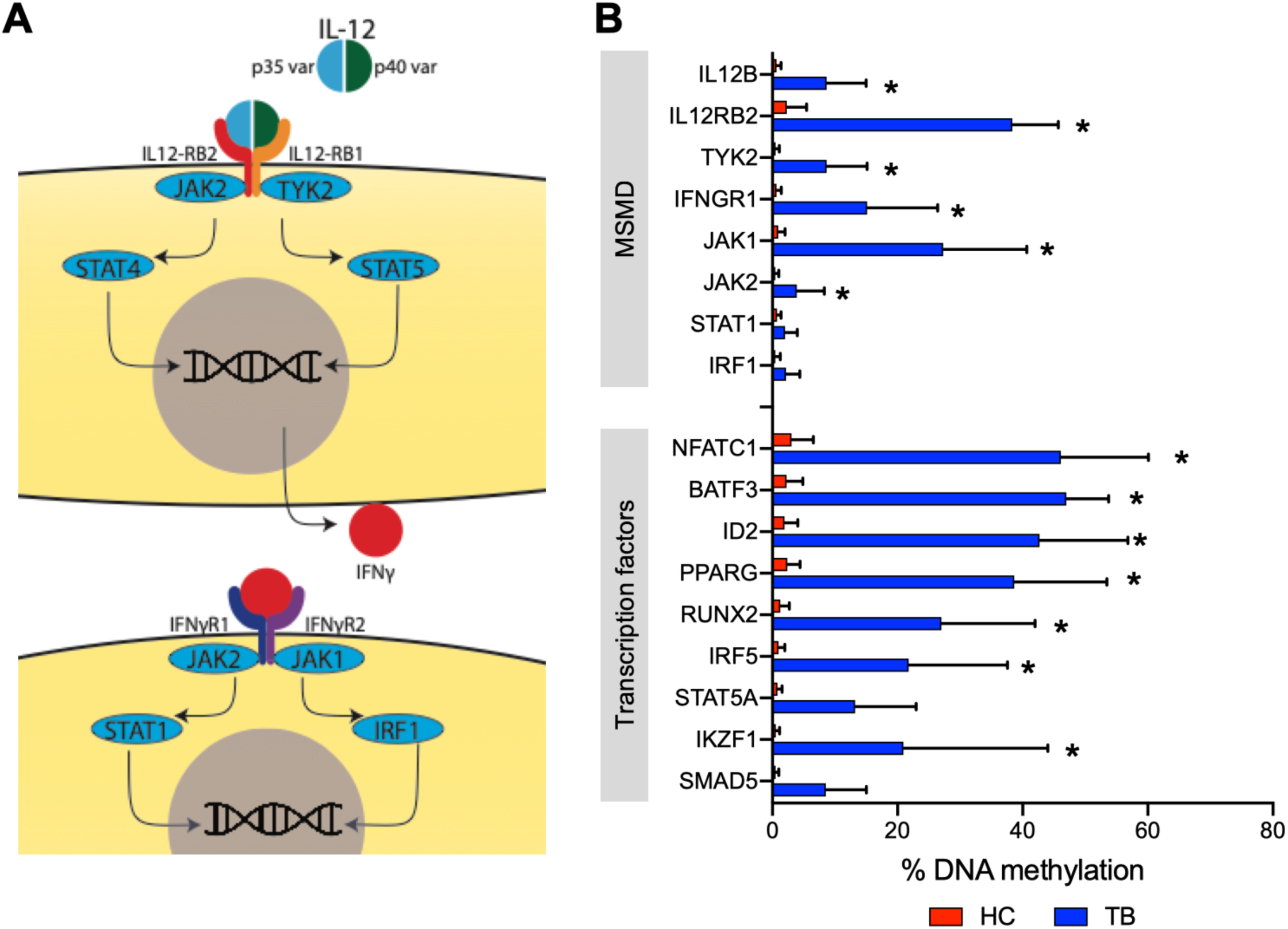
Targeted DNA methylation confirms DNA hyper-methylation of the IL-12-IFN-γ pathway. (**A**) Graphic representation of the IL-12-IFN-γ pathway (**B**) DNA methylation was evaluated using ***m***ethylation ***s***pecific ***r***estriction ***e***ndonuclease isochizomers and PCR (MSRE-qPCR) in non-HIV co-infected TB patients (n = 6) and their asymptomatic healthy household contacts (n = 5). Mann-Whitney p-value < 0.01 for all genes depicted by *.

### TB patients have decreased mitogen responsiveness

TB patients demonstrated DNA hyper-methylation of IL-2, IL-6, TNF and IFN-γ signaling pathways (Fig 2-4). Each of these pathways result in activation of mitogen activated protein kinase (MAPK) and each cytokine is also downstream of mitogen activation. We therefore hypothesized that TB patients would have decreased mitogen responsiveness. Freshly collected whole blood was stimulated overnight with mitogen and multiplexed ELISA was performed on the supernatant. The TNF-NFϰB signaling pathway and the gene *TNF* were hyper-methylated in CD14, CD8 and CD56 cells of TB patients (Fig. 3). While the baseline production of TNF was similar (3.7 vs 1.4 pg/mL; p = 0.2), after mitogen stimulation, TB patients exhibited 72% decreased up regulation of TNF by ELISA (80.5 vs 293 pg/ mL; p = 0.01; Fig. 5). The IFN-γ signaling pathway was hyper-methylated in all lymphocytes and the *IFNG* gene was hyper-methylated in CD8 and CD56 cells from TB patients and while TB patients had increased IFN-γ by ELISA at baseline (1.31 vs 0.58 pg/mL, p = 0.007), they demonstrated 35.4% decreased capacity to upregulate IFN-γ upon mitogen activation (increase 246.9 vs 382.4 pg/mL; p = 0.03). IL-6 and IL-1β levels were comparable at baseline, but TB patients had 40.5 and 60.7% decreased mitogen-induced upregulation of IL-6 (8870 vs 14917 pg/mL; p < 0.001) and IL-1β (1968 vs 5974 pg/mL, p = 0.0005), respectively. The chemokines CXCL9 and CXCL10, which are downstream of both IL12 and IFN-γ signaling, were increased at baseline (42.5 vs 10.6 and 238.3 vs. 117.4 pg/ml; p < 0.001), but not after mitogen induced activation.

**Figure 5:**
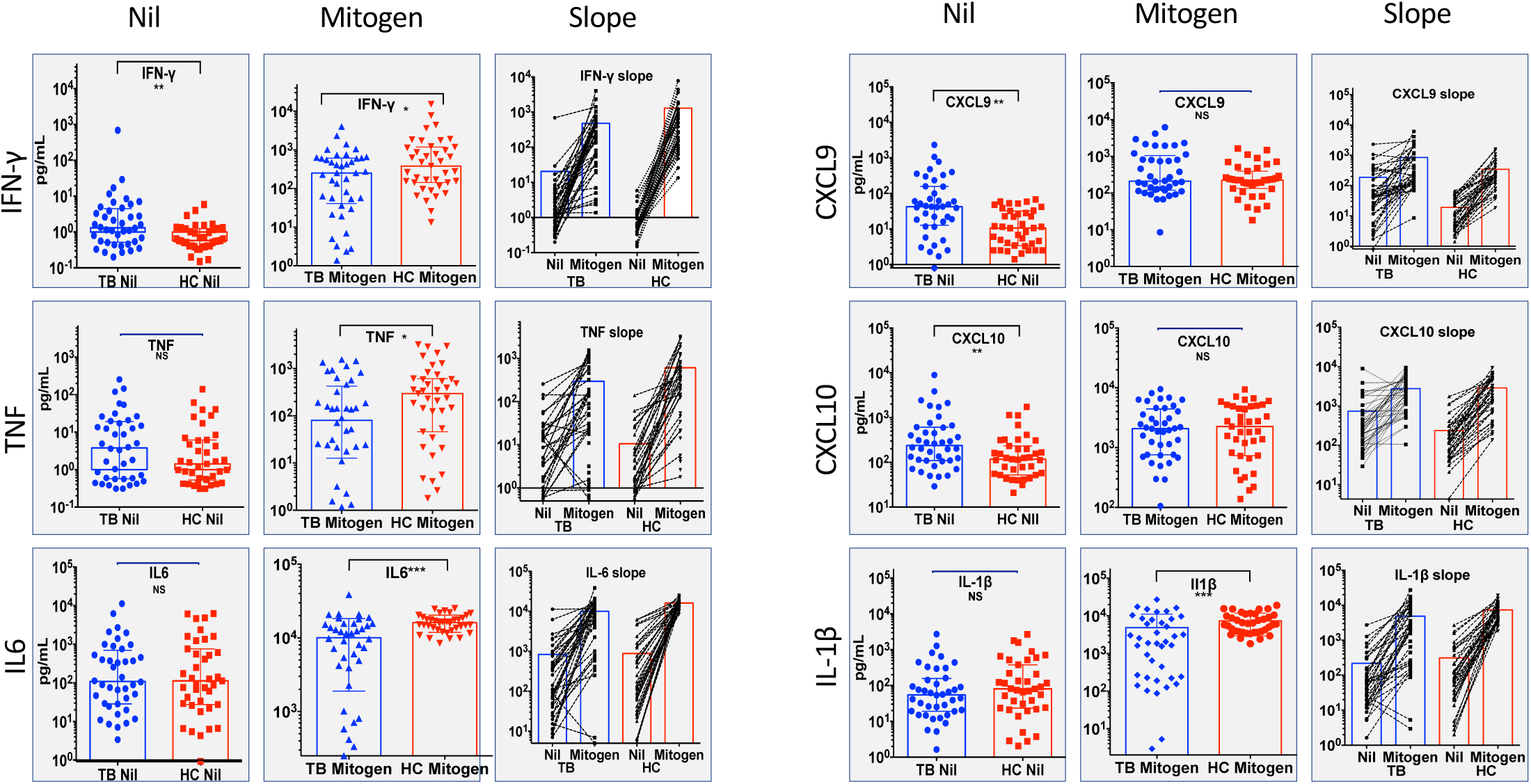
Decreased mitogen-induced responsiveness. Fresh whole blood (0.8 – 1.2 mL) from TB patients (n = 40) and asymptomatic healthy household contacts (n = 39) was stimulated overnight with and without mitogen and the resultant plasma was evaluated for cytokine and chemokine upregulation using a bead-based multiplex ELISA. Mann-Whitney p-values < 0.05, < 0.01 and < 0.001 depicted by *, **, and *** respectively.

### TB patients have decreased IFN-γ induced gene upregulation

An intact IL-12-IFN-γ pathway is necessary, but not sufficient, for control of mycobacterial infections^25,26^. Since TB patients exhibited DNA hyper-methylation of the IL-12-IFN-γ signaling pathway in CD4^+^ helper T cells, NK cells and CTLs, we postulated they would have decreased IFN-γ inducible gene upregulation using a previously described ten-gene IFN-γ inducible score^14,27,28^. In healthy immune cells, IFN-γ stimulation induces gene expression of *CITTA, GBP1, STAT1, CXCL9, CXCL10, CXCL11, IL15, SERPING1, IDO1, FCGR1A/B*^*27*^, however previous studies have demonstrated decreased IFN-γ-inducible gene expression in cancer^28^ and chronic schistosomiasis infection^14^. In this cohort, compared to controls, the PBMCs from TB patients demonstrated 69.7% decreased IFN-γ-inducible gene upregulation (17.9 vs 59.2 fold-increase; p = 0.02; Fig. 6).

**Figure 6:**
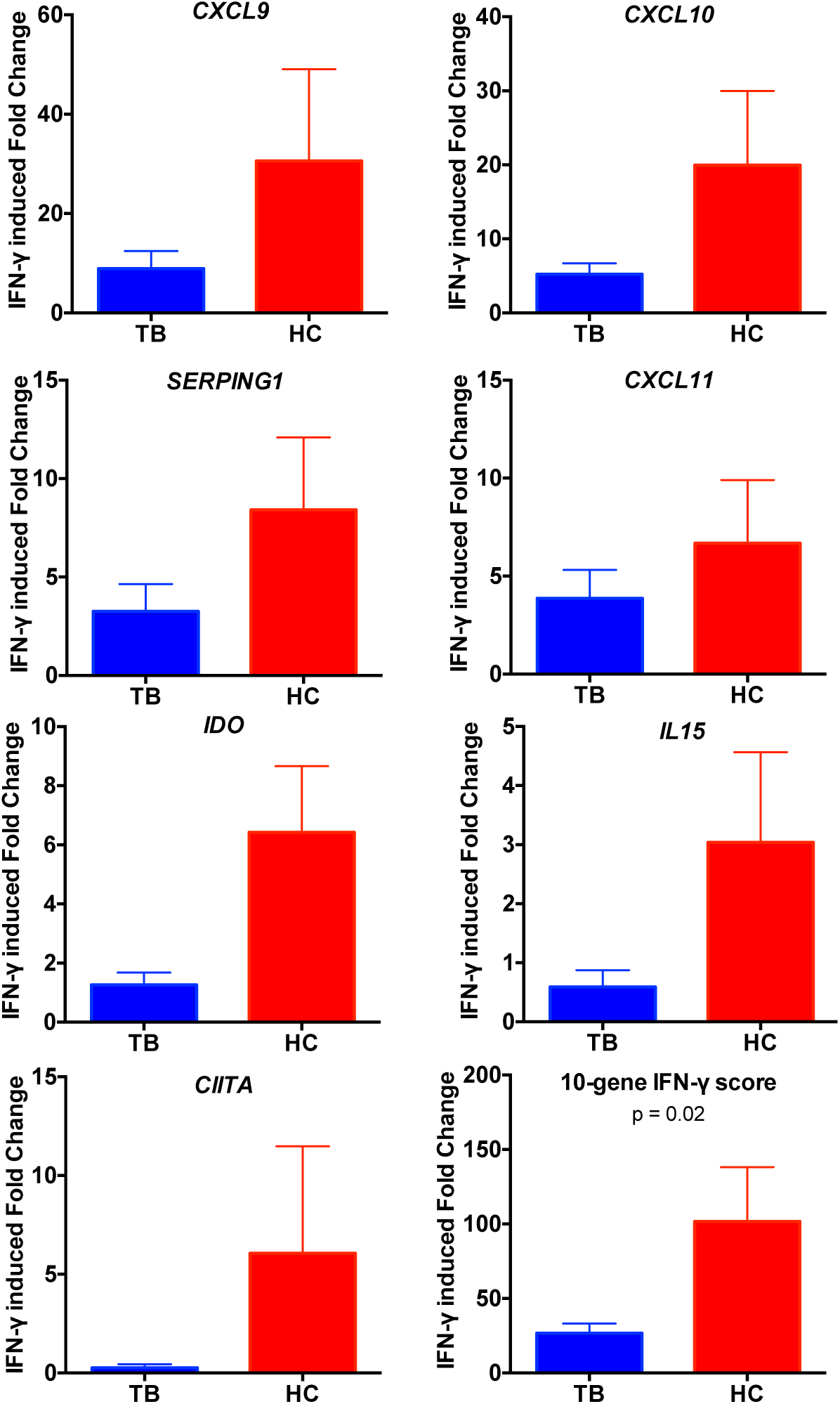
Decreased IFN-γ inducible gene expression. (**A**) PBMCs (1×10^6^) from TB patients (n = 10) and asymptomatic healthy household contacts (n = 10) was stimulated overnight with and without 50ng of IFN-γ. The following morning RNA was isolated and IFN-γ-inducible gene expression evaluated by microarray.

### TB patients have decreased IL-12 inducible IFN-γ production

Lymphocytes are primed with IL-12 for robust IFN-γ production^29^. Since TB patients demonstrated DNA hyper-methylation of the IL12 pathway (*IL12A, IL12B, IL12RB1 and IL12RB2*), we evaluated the functionality of the IL-12-IFN-γ axis by stimulating PBMCs overnight with BCG with and without IL-12. While helper T cell (CD3^+^CD4^+^) IFN-γ production was similar, TB patients demonstrated a muted IL-12 induced production of IFN-γ in CTLs and NK cells. Specifically, NK cells had 62.5% decreased IL-12-inducible up-regulation of IFN-γ (p < 0.01), while CTLs had a 43.4% decrease in IL-12-induced IFN-γ up-regulation (p = 0.03; Fig.7).

**Figure 7:**
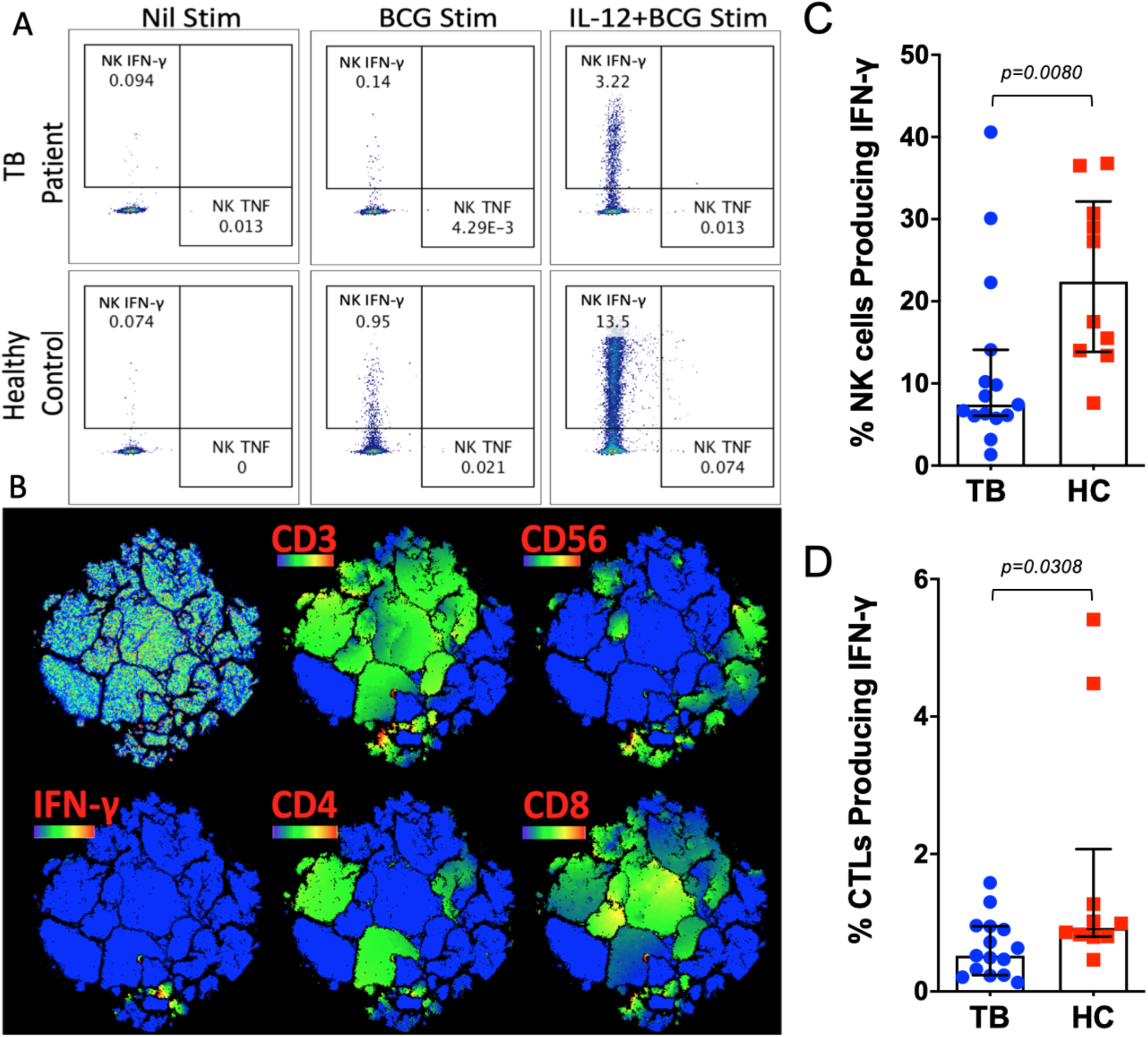
Decreased IL-12 inducible IFN-γ production. PBMCs from TB patients (n = 15) and asymptomatic healthy household contacts (n = 10) were stimulated overnight without stimulation (PBS media control), with BCG sonicate (5μg) or with IL-12 (20ng/mL) and BCG (minimum of 0.5 x10^6^ PBMCs per condition) and evaluated by multi-dimensional flow cytometry the following morning. (**A**) Representative dot plots of NK (CD3^-^CD56^+^) production of IFN-γ and TNFα after no stimulation (“Nil stim”), BCG, or IL-12 and BCG stimulation. (**B**) tSNE (t-Distributed Stochastic Neighbor Embedding dimension reduction) clustering demonstrating the cell-specific (CD3, CD4, CD8 and CD56) contribution of IFN-γ production after IL-12 and BCG stimulation with blue indicating no protein expression, and green, yellow and red indicating increased protein expression. (**C-D**) Bar graphs demonstrating the percent of CTL (CD3^+^CD8^+^ and NK cell (CD3^-^CD56^+^) producing IFN-γ after IL-12 and BCG stimulation. Mann-Whitney p = 0.3 and 0.008 for CTLs and NK cells, respectively.

## Discussion

Despite becoming culture-negative after two months of anti-tuberculosis therapy, patients with TB require six grueling months of antibiotics^30^. However, even after these six months of antibiotics, patients with successful anti-TB therapy have a 13-fold increased risk of recurrent TB compared to the general population^2^. Epigenetic-mediated immune exhaustion has been demonstrated to impede immunotherapy in murine LCMV experimental models^9^, but it has not previously been evaluated as a potential etiology of TB induced immune exhaustion. Here we demonstrate that TB patients have DNA hyper-methylation in critical immune pathways including the IL-12-IFN-γ pathway and this is associated with decreased mitogen, mycobacterial, IL-12 and IFN-γ immune responsiveness.

Anergy in TB patients has been described since at least the 1940s. Depending on the definition employed for anergy, approximately 5 - 25% of TB patients fail to mount a positive skin reaction to tuberculin, or failure to release IFN-γ after exposure to mycobacterial antigens^31-37^. TB patients failing to produce *Mtb*-specific IFN-γ have increased mortality^38^, potentially suggesting more advanced disease. The TB-induced anergy is not specific to mycobacterial antigens, with TB patients also exhibiting decreased responsiveness to *Candida* and histoplasmin antigens^39^. HIV increases the likelihood of anergy^34^, but in some cohorts up to 46% of HIV-negative TB cases are anergic^40^. Modern immune analysis has characterized TB-induced immune exhaustion demonstrating that TB patients have decreased IFN-γ, IL-2 and proliferative capacity with an increase in immune checkpoint inhibitors, such as PD-1, TIM3 and CTLA-4^18,19,41,42^. In addition to limiting critical cytokines, such as IFN-γ, previous studies have demonstrated that immune cells from TB patients to have downstream defects in cell signaling including decreased IFN-γ-inducible gene expression^43,44^. DNA hyper-methylation has been demonstrated to inhibit robust immunity in LCMV models, and here we identify that the DNA hyper-methylation is associated with suppressed immunity during TB.

Immune exhaustion after chronic antigen stimulation is not limited to TB and has been observed in many instances of persistent inflammation due to chronic infection, sepsis or cancer^6,45-47^. Continuously expressed tumor antigens induce immune exhaustion via upregulation of DNA methyltransferases and the transcriptionally repressive EZH2 complex, both known moderators of epigenetic-mediated immune exhaustion^12,48^. Epigenetic-mediated immune exhaustion persists even if cells are rested for three weeks, or if they are transferred into healthy, non-infected mice^10,12^. In clinical cohorts, even two years after successful aviremia, patients infected with HIV retain detrimental epigenetic marks^8,13^. Similarly, 6 months after successful deworming, children previously schistosomiasis infected retain detrimental DNA methylation marks^14^. In this report, six-months after completing successful anti-TB therapy, the immune system demonstrates persistent DNA hyper-methylation perturbations. Patients in this cohort were only followed six months after completion of ATT, and longitudinal studies will need to assess if persistent detrimental DNA methylation marks are correlative with recurrent TB disease.

Chronic LCMV models have demonstrated that immune exhaustion is moderated by broad remodeling of the epigenetic landscape and that these epigenetic perturbations encumber immune checkpoint blockade-based immunotherapy^4,5^. Ghoneim et al demonstrated that immune exhaustion is moderated by DNA methylation with hypomethylating agents dosed prior to immune checkpoint inhibitor therapy capable of reversing epigenetic-mediated immune exhaustion^9^. This data demonstrates that the DNA hyper-methylation landscape of TB patients before and after successful anti-TB therapy resembles murine LCMV-induced immune exhaustion regions of closed chromatin by chronic LCMV (Sen et al) and DNA hyper-methylation (Ghoneim et al). In addition to chronic antigen induced epigenetic-mediated immune exhaustion, an alternative and non-mutually exclusive mechanism of host epigenetic perturbations is pathogen production of host epigenetic-modifying enzymes. Leishmania, HIV, HPV and *Mtb* all produce epigenetic-modifying enzymes that target host epigenetic status^22,23,44,49-53^. *Mtb* itself produces a functional DNA methyltransferase that is secreted, chaperoned into the nucleus, and induces host DNA methylation changes in the IL-12-IFN-γ signaling^22,23^ similar to the DNA methylation changes demonstrated in this cohort (Supplemental Figure 2). The current study did not evaluate the *Mtb* strain for DNMT production, but future studies should evaluate the effect of strain on host DNA methylation in clinical cohorts.

The DNA hyper-methylation of the IL-12-IFN-γ signaling pathway, especially among CD8 and NK cell populations, may partially explain the complexity of identifying IFN-γ mediated immune correlates of protection. The Mendelian Susceptibility to Mycobacterial Diseases (MSMDs) are a rare collection of gene mutations that increase an individual’s risk for Tuberculosis and other intracellular infections. The MSMDs highlights the critical nature and complexity of this pathway. Upstream mutations in *IL12B, IL12RB*, or *NEMO* result in decreased IFN-γ, decreased IFN-γ-inducible gene expression and decreased mycobacterial killing capacity. In contrast, mutations downstream of IFN-γ in *IFNGR1, IFNGR2*, or *STAT1* result in increased IFN-γ, but decreased IFN-γ-inducible gene expression and decreased mycobacterial killing capacity. Here, we demonstrate that TB patients have increased DNA methylation in the canonical Il-12-IFN-γ pathway (Fig 4a) both upstream (in *IL12, IL12RB2, STAT4)* and downstream (in *IFNG, JAK1* and *STAT1)* of IFN-γ (Fig. 3-4). However, IFN-γ gene expression response is dependent upon both canonical and non-canonical signaling. Non-canonical modulators of IFN-γ-inducible gene expression include *FYN, MAL* and the mTOR pathway, which were also hyper-methylated in TB patients. DNA hyper-methylation of the canonical and non-canonical IL-12-IFN-γ pathway was associated with reduced IFN-γ inducible gene expression (Fig. 6), reduced mitogen-induced cytokine production (Fig. 5) and reduced IL-12-inducible IFN-γ production (Fig. 7).

Host-directed immunotherapy has been identified as a priority research area to improve TB treatment outcomes among the 10 million people afflicted each year. TB is an archetypical chronic infection known to inhibit host immunity and herein, we demonstrate that decreased immune responsiveness to IL-12, IFN-γ, mitogen, and mycobacterial antigens is associated with DNA hyper-methylation of critical immune transcription factors and the canonical and non-canonical IL-12-IFN-γ signaling pathway. The persistent DNA methylation perturbations that we have demonstrated are a plausible explanation for why successful TB therapy must be continued for months after *Mtb* culture conversion. These data suggest it is critical to now evaluate whether modulating DNA methylation can effectively augment host antimycobacterial immunity.

## Methods

### Ethics, cohort enrollment and study samples

The study protocol was reviewed by the Baylor College of Medicine Children’s Foundation-Swaziland, Baylor College of Medicine and Eswatini National Health Research and Review Board. All participation was voluntary and implemented in accord with the institutional and international guidelines for the Protection of Human Subjects in concordance with the Declaration of Helsinki. In this analysis, immune cells of TB participants were analyzed if they had pulmonary disease as defined by symptoms of TB (cough, fevers, weight loss, etc) and microbiologically confirmed disease by sputum Gene Xpert and/ or culture. All participants were screened for HIV. Individuals with helminth co-infection (by urine or stool microscopy, and urine and stool qPCR) were excluded from this analysis. Healthy controls were household contacts of TB patients with a TB contact score^54^ of five or greater and excluded if they progressed to active TB and did not remain asymptomatic for 12-months. All individuals were BCG vaccinated as confirmed by BCG scar and/ or vaccine card.

### Multi-Dimensional Immune Profiling

Peripheral blood mononuclear cells were isolated using Ficoll-separation and cryopreserved in liquid nitrogen. Upon thawing, >70% viability was determined by trypan and by lymphocyte amine reactive dye by flow cytometry. PBMCs were stimulated under the following conditions: a) DMSO vehicular control, b) 2.5 μg each of ESAT-6 and CFP-10 overlapping peptide pools, c) 5 μg of BCG sonicate and d) 0.2 μL of staphylococcal enterotoxin B. Stimulations took place in the presence of co-stimulation (CD28/CD49) for 18 hours with monensin and brefeldin A for the last 12 hours. Cells were stained with an amine reactive dye (Ghost Dye), surface antibodies (CD4, CD8, CD56 and PD-1) and intracellular antibodies (CD3, Ki-67, IFN-γ, TNF, IL-4, IL-10, IL-13, perforin, T-bet and GATA-3) and acquired on an LSR II Fortessa.

### DNA methylation

Genomic DNA was isolated using the DNeasy kit from 1×10^6^ PBMC or magnetic-bead isolated CD4^+^ T cells with a minimum of 80% viability. Nucleic acid quantity and quality was examined using a Qubit 3.0 fluorometer and Agilent 4200 Tape Station System and 500ng of gDNA was bisulfite treated prior to running the Illumina Infinium MethylEPIC array. Methylation IDAT files were preprocessed and normalized using the R statistical system Bioconductor) minfi package with probes with greater than 2, or less than 0.5-fold differential methylation with p-values < 0.05 being considered significant. Gene ontology (GO) and Gene Set Enrichment Analysis was implemented using the Molecular Signature Database (MSigDB) using hyper-geometric distribution that accounts for multiple comparisons.^55^ Epigenetic deconvolution was implemented as previously described^56^. The Epitect II DNA Methylation Enzyme kit was used to validate methylome results as previously described. In brief, gDNA underwent methylation specific enzymatic digestion with the percentage of methylated and unmethylated DNA determined relative to mock and double digest determined by quantitative PCR^14,57^.

### Gene expression

PBMCs were stimulated overnight with IFN-γ (50 ng/mL) or media control (PBS) for 16 hours followed by RNA isolation and gene expression evaluated using the Nanostring nCounter Human Immunology 2 probe set. Data was normalized using eight negative controls and ten housekeeping genes.

### Statistical analysis

Flow cytometry data was processed using FlowJo (TreeStar, Ashland, OR) and GraphPad 6.0 (graphPad Software). CITRUS was implemented using CytoBank (http://www.cytobank.org) using nearest shrunken centroid (PAMR) with a minimum cluster size of 10%, cross validation fold of 10 and an FDR of 1%. DNA methylEPIC probes were considered significant if they demonstrated greater than 2 or less than 0.5-fold differences from controls with a p-value < 0.05. Gene expression, flow cytometry and targeted MSRE DNA methylation was evaluated using nonparametric Mann-Whitney U test.

## Figures

**Supplemental Figure 1:**
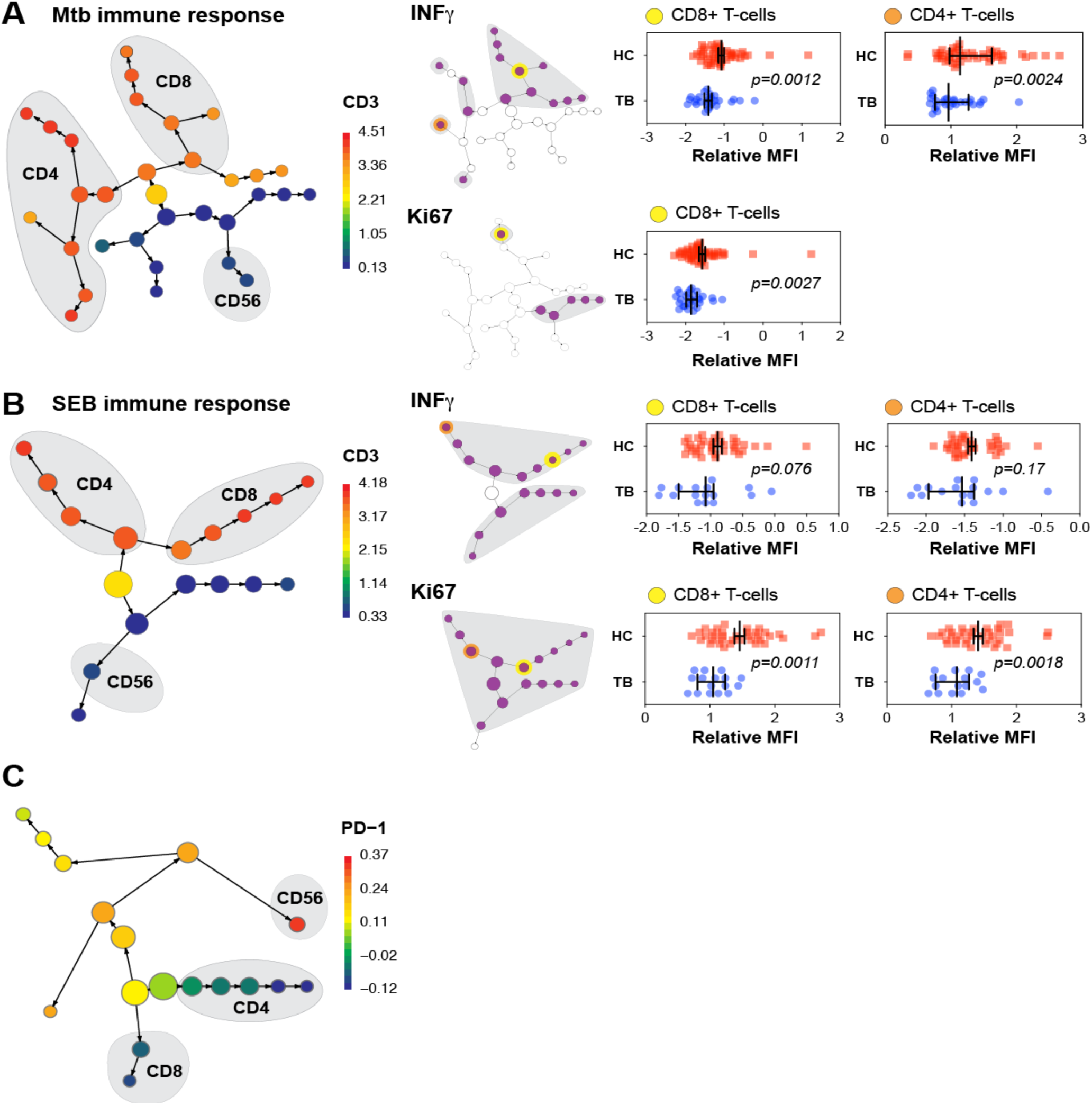
TB patients exhibit an exhausted immune phenotype. PBMCs from individuals with Tuberculosis (TB; n = 29) and their asymptomatic household contacts (HC; n = 49) had their PBMCs divided into PBS vehicle non-stimulated control (“Nil stim”), BCG stimulated, ESAT-6 and CFP-10 stimulated and Staphylococcal-enterotoxin B (SEB) stimulated cells (0.5 – 1 x 10^6^ cells per condition). Cells were stimulated overnight with monensin and BFA for the final 12 hours. The BCG immune response is presented in Figure 1. (**S1A**) Citrus (cluster identification, characterization, and regression) identified CD4, CD8, CD3 and CD56 subsets. (**S1A**) Mtb-specific immune response: CITRUS clustering identifies CD4, CD8 and NK populations with CD3 signal depicted by color gradient. After stimulation with ESAT-6 and CFP-10, *Mtb*-specific peptides, TB participants demonstrate decreased IFN-γ production by CD4 and CD8 T cells. (**S1B**) Staphylococcal-enterotoxin B (SEB) immune response: CITRUS analysis demonstrates CD4, CD8 and NK cells with decreased Ki-67 and IFN-γ. (**S1C**) PD-1 NK cells: CITRUS analysis identifies TB patients to have increased frequency of PD-1 expressing NK cells compared to healthy controls.

**Supplemental Figure 2:**
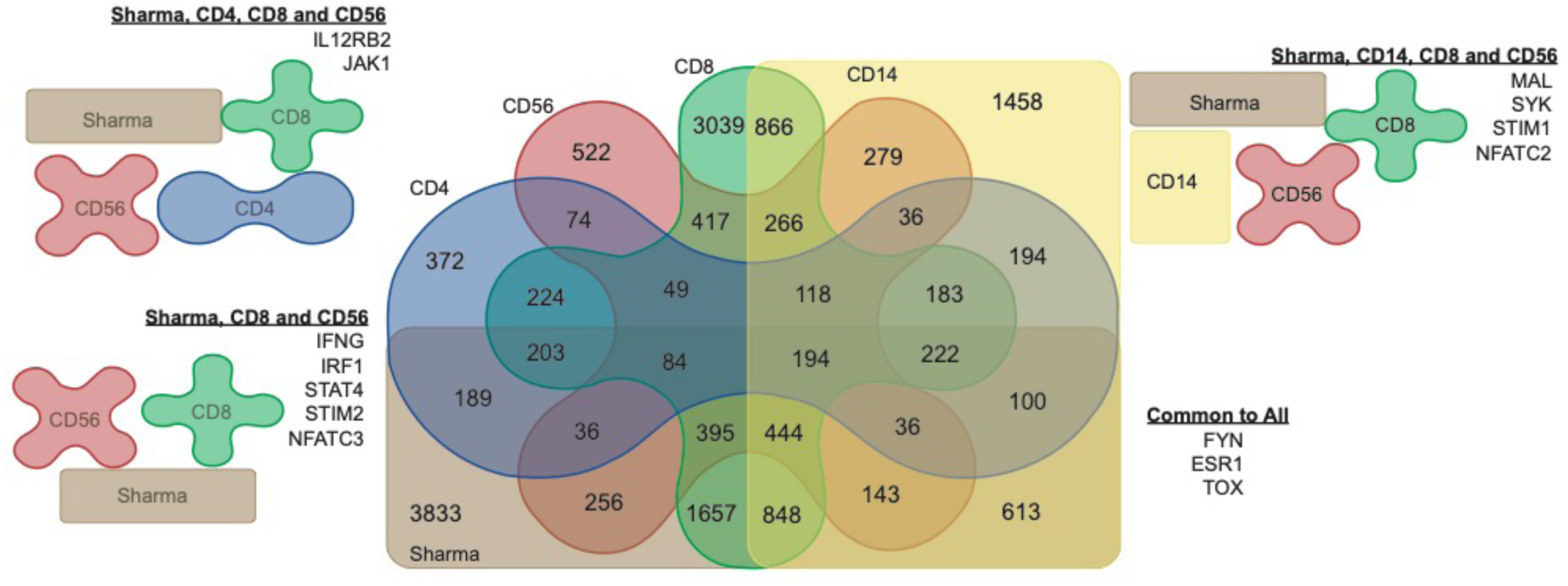
Venn Diagram demonstrates overlap of in vivo DNA hyper-methylation in TB patients from Eswatini with previously published in vitro DNA hyper-methylation (Sharma et al). Venn diagram of genes with DNA hyper-methylation in CD4, CD8, CD56 and CD14 cells from this cohort of TB patients.

## Author Contributions

ARD and AMM designed the study and wrote the manuscript. GM, GM, AK, ARD, QD, and JK implemented cohort enrollment and clinical study design. TN, ARD, QD and JK implemented the immunology and DNA methylation. SM, SG and ARD implemented CITRUS analysis. KR, SG and CC implemented the bioinformatic and statistical analysis. GM, EM, JDC, MGN, and RvC assisted with data interpretation and designing biologic validation studies.

## Acknowledgements

This work was supported by Burroughs Wellcome Fund/ American Society of Tropical Medicine and Hygiene–Postdoctoral Fellowship in Tropical Infectious Diseases (early career grant), Thrasher Research Fund (early career grant), and the National Institutes of Health (K23AI141681A).

The authors have declared no conflicts of interest exist

## REFERENCES

1. WHO. Global tuberculosis report. World Health Organization, Geneva, Switzerland. 2016.

2. Millet JP, Shaw E, Orcau A, Casals M, Miro JM, Cayla JA, Barcelona Tuberculosis Recurrence Working G. Tuberculosis recurrence after completion treatment in a European city: reinfection or relapse? PLoS One 2013;8(6):e64898.

3. Ahmed R, Salmi A, Butler LD, Chiller JM, Oldstone MB. Selection of genetic variants of lymphocytic choriomeningitis virus in spleens of persistently infected mice. Role in suppression of cytotoxic T lymphocyte response and viral persistence. J Exp Med 1984;160(2):521–40.

4. Sen DR, Kaminski J, Barnitz RA, Kurachi M, Gerdemann U, Yates KB, Tsao HW, Godec J, LaFleur MW, Brown FD, et al. The epigenetic landscape of T cell exhaustion. Science 2016;354(6316):1165–9.

5. Pauken KE, Sammons MA, Odorizzi PM, Manne S, Godec J, Khan O, Drake AM, Chen Z, Sen DR, Kurachi M, et al. Epigenetic stability of exhausted T cells limits durability of reinvigoration by PD-1 blockade. Science 2016;354(6316):1160–5.

6. Martinez GJ, Pereira RM, Aijo T, Kim EY, Marangoni F, Pipkin ME, Togher S, Heissmeyer V, Zhang YC, Crotty S, et al. The transcription factor NFAT promotes exhaustion of activated CD8(+) T cells. Immunity 2015;42(2):265–78.

7. Pauken KE, Wherry EJ. SnapShot: T Cell Exhaustion. Cell 2015;163(4):1038–e1.

8. Nakayama-Hosoya K, Ishida T, Youngblood B, Nakamura H, Hosoya N, Koga M, Koibuchi T, Iwamoto A, Kawana-Tachikawa A. Epigenetic repression of interleukin 2 expression in senescent CD4+ T cells during chronic HIV type 1 infection. J Infect Dis 2015;211(1):28–39.

9. Ghoneim HE, Fan Y, Moustaki A, Abdelsamed HA, Dash P, Dogra P, Carter R, Awad W, Neale G, Thomas PG, et al. De Novo Epigenetic Programs Inhibit PD-1 Blockade-Mediated T Cell Rejuvenation. Cell 2017;170(1):142–57 e19.

10. Ahn E, Youngblood B, Lee J, Lee J, Sarkar S, Ahmed R. Demethylation of the PD-1 Promoter Is Imprinted during the Effector Phase of CD8 T Cell Exhaustion. J Virol 2016;90(19):8934–46.

11. Miller BC, Sen DR, Al Abosy R, Bi K, Virkud YV, LaFleur MW, Yates KB, Lako A, Felt K, Naik GS, et al. Subsets of exhausted CD8(+) T cells differentially mediate tumor control and respond to checkpoint blockade. Nat Immunol 2019;20(3):326–36.

12. Schietinger A, Philip M, Krisnawan VE, Chiu EY, Delrow JJ, Basom RS, Lauer P, Brockstedt DG, Knoblaugh SE, Hammerling GJ, et al. Tumor-Specific T Cell Dysfunction Is a Dynamic Antigen-Driven Differentiation Program Initiated Early during Tumorigenesis. Immunity 2016;45(2):389–401.

13. Youngblood B, Noto A, Porichis F, Akondy RS, Ndhlovu ZM, Austin JW, Bordi R, Procopio FA, Miura T, Allen TM, et al. Cutting edge: Prolonged exposure to HIV reinforces a poised epigenetic program for PD-1 expression in virus-specific CD8 T cells. J Immunol 2013;191(2):540–4.

14. DiNardo AR, Nishiguchi T, Mace EM, Rajapakshe K, Mtetwa G, Kay A, Maphalala G, Secor WE, Mejia R, Orange JS, et al. Schistosomiasis Induces Persistent DNA Methylation and Tuberculosis-Specific Immune Changes. J Immunol 2018.

15. Ringeard M, Marchand V, Decroly E, Motorin Y, Bennasser Y. FTSJ3 is an RNA 2’-O-methyltransferase recruited by HIV to avoid innate immune sensing. Nature 2019;565(7740):500–4.

16. Gunther G, Lange C, Alexandru S, Altet N, Avsar K, Bang D, Barbuta R, Bothamley G, Ciobanu A, Crudu V, et al. Treatment Outcomes in Multidrug-Resistant Tuberculosis. N Engl J Med 2016;375(11):1103–5.

17. Bruggner RV, Bodenmiller B, Dill DL, Tibshirani RJ, Nolan GP. Automated identification of stratifying signatures in cellular subpopulations. Proc Natl Acad Sci U S A 2014;111(26):E2770–7.

18. Singh A, Mohan A, Dey AB, Mitra DK. Inhibiting the programmed death 1 pathway rescues Mycobacterium tuberculosis-specific interferon gamma-producing T cells from apoptosis in patients with pulmonary tuberculosis. J Infect Dis 2013;208(4):603–15.

19. Day CL, Abrahams DA, Bunjun R, Stone L, de Kock M, Walzl G, Wilkinson RJ, Burgers WA, Hanekom WA. PD-1 Expression on Mycobacterium tuberculosis-Specific CD4 T Cells Is Associated With Bacterial Load in Human Tuberculosis. Front Immunol 2018;9:1995.

20. Roy Chowdhury R, Vallania F, Yang Q, Lopez Angel CJ, Darboe F, Penn-Nicholson A, Rozot V, Nemes E, Malherbe ST, Ronacher K, et al. A multi-cohort study of the immune factors associated with M. tuberculosis infection outcomes. Nature 2018;560(7720):644–8.

21. Onuchic V, Hartmaier RJ, Boone DN, Samuels ML, Patel RY, White WM, Garovic VD, Oesterreich S, Roth ME, Lee AV, et al. Epigenomic Deconvolution of Breast Tumors Reveals Metabolic Coupling between Constituent Cell Types. Cell Rep 2016;17(8):2075–86.

22. Sharma G, Upadhyay S, Srilalitha M, Nandicoori VK, Khosla S. The interaction of mycobacterial protein Rv2966c with host chromatin is mediated through non-CpG methylation and histone H3/H4 binding. Nucleic Acids Res 2015;43(8):3922–37.

23. Sharma G, Sowpati DT, Singh P, Khan MZ, Ganji R, Upadhyay S, Banerjee S, Nandicoori VK, Khosla S. Genome-wide non-CpG methylation of the host genome during M. tuberculosis infection. Sci Rep 2016;6:25006.

24. Evans CM, Jenner RG. Transcription factor interplay in T helper cell differentiation. Brief Funct Genomics 2013;12(6):499–511.

25. Orange JS, Levy O, Brodeur SR, Krzewski K, Roy RM, Niemela JE, Fleisher TA, Bonilla FA, Geha RS. Human nuclear factor kappa B essential modulator mutation can result in immunodeficiency without ectodermal dysplasia. J Allergy Clin Immunol 2004;114(3):650–6.

26. Bustamante J, Boisson-Dupuis S, Abel L, Casanova JL. Mendelian susceptibility to mycobacterial disease: genetic, immunological, and clinical features of inborn errors of IFN-gamma immunity. Semin Immunol 2014;26(6):454–70.

27. Waddell SJ, Popper SJ, Rubins KH, Griffiths MJ, Brown PO, Levin M, Relman DA. Dissecting interferon-induced transcriptional programs in human peripheral blood cells. PLoS One 2010;5(3):e9753.

28. Ayers M, Lunceford J, Nebozhyn M, Murphy E, Loboda A, Kaufman DR, Albright A, Cheng JD, Kang SP, Shankaran V, et al. IFN-gamma-related mRNA profile predicts clinical response to PD-1 blockade. J Clin Invest 2017;127(8):2930–40.

29. Wysocka M, Kubin M, Vieira LQ, Ozmen L, Garotta G, Scott P, Trinchieri G. Interleukin-12 is required for interferon-gamma production and lethality in lipopolysaccharide-induced shock in mice. Eur J Immunol 1995;25(3):672–6.

30. Gillespie SH, Crook AM, McHugh TD, Mendel CM, Meredith SK, Murray SR, Pappas F, Phillips PP, Nunn AJ, Consortium RE. Four-month moxifloxacin-based regimens for drug-sensitive tuberculosis. N Engl J Med 2014;371(17):1577–87.

31. Zeitz SJ, Ostrow JH, Van Arsdel PP, Jr. Humoral and cellular immunity in the anergic tuberculosis patient. A prospective study. J Allergy Clin Immunol 1974;53(1):20–6.

32. Holden M, Dubin MR, Diamond PH. Frequency of negative intermediate-strength tuberculin sensitivity in patients with active tuberculosis. N Engl J Med 1971;285(27):1506–9.

33. Michael L. Furcolow BH, Waldo E. Nelson and Carroll E. Palmer. Quantitative Studies of the Tuberculin Reaction: I. Titration of Tuberculin Sensitivity and Its Relation to Tuberculous Infection. Public Health Reports 1941;Vol. 56, No. 21:1082–100.

34. Cobelens FG, Egwaga SM, van Ginkel T, Muwinge H, Matee MI, Borgdorff MW. Tuberculin skin testing in patients with HIV infection: limited benefit of reduced cutoff values. Clin Infect Dis 2006;43(5):634–9.

35. Nash DR, Douglass JE. Anergy in active pulmonary tuberculosis. A comparison between positive and negative reactors and an evaluation of 5 TU and 250 TU skin test doses. Chest 1980;77(1):32–7.

36. Sollai S, Galli L, de Martino M, Chiappini E. Systematic review and meta-analysis on the utility of Interferon-gamma release assays for the diagnosis of Mycobacterium tuberculosis infection in children: a 2013 update. BMC Infect Dis 2014;14 Suppl 1:S6.

37. Metcalfe JZ, Everett CK, Steingart KR, Cattamanchi A, Huang L, Hopewell PC, Pai M. Interferon-gamma release assays for active pulmonary tuberculosis diagnosis in adults in low- and middle-income countries: systematic review and meta-analysis. J Infect Dis 2011;204 Suppl 4:S1120–9.

38. Nguyen DT, Teeter LD, Graves J, Graviss EA. Characteristics Associated with Negative Interferon-gamma Release Assay Results in Culture-Confirmed Tuberculosis Patients, Texas, USA, 2013-2015. Emerg Infect Dis 2018;24(3):534–40.

39. McMurray DN, Echeverri A. Cell-mediated immunity in anergic patients with pulmonary tuberculosis. Am Rev Respir Dis 1978;118(5):827–34.

40. Sousa AO, Salem JI, Lee FK, Vercosa MC, Cruaud P, Bloom BR, Lagrange PH, David HL. An epidemic of tuberculosis with a high rate of tuberculin anergy among a population previously unexposed to tuberculosis, the Yanomami Indians of the Brazilian Amazon. Proc Natl Acad Sci U S A 1997;94(24):13227–32.

41. Sahiratmadja E, Alisjahbana B, de Boer T, Adnan I, Maya A, Danusantoso H, Nelwan RH, Marzuki S, van der Meer JW, van Crevel R, et al. Dynamic changes in pro- and anti-inflammatory cytokine profiles and gamma interferon receptor signaling integrity correlate with tuberculosis disease activity and response to curative treatment. Infect Immun 2007;75(2):820–9.

42. Harari A, Rozot V, Enders FB, Perreau M, Stalder JM, Nicod LP, Cavassini M, Calandra T, Blanchet CL, Jaton K, et al. Dominant TNF-alpha+ Mycobacterium tuberculosis-specific CD4+ T cell responses discriminate between latent infection and active disease. Nat Med 2011;17(3):372–6.

43. Kincaid EZ, Ernst JD. Mycobacterium tuberculosis exerts gene-selective inhibition of transcriptional responses to IFN-gamma without inhibiting STAT1 function. J Immunol 2003;171(4):2042–9.

44. Pennini ME, Liu Y, Yang J, Croniger CM, Boom WH, Harding CV. CCAAT/enhancer-binding protein beta and delta binding to CIITA promoters is associated with the inhibition of CIITA expression in response to Mycobacterium tuberculosis 19-kDa lipoprotein. J Immunol 2007;179(10):6910–8.

45. Doering TA, Crawford A, Angelosanto JM, Paley MA, Ziegler CG, Wherry EJ. Network analysis reveals centrally connected genes and pathways involved in CD8+ T cell exhaustion versus memory. Immunity 2012;37(6):1130–44.

46. Mognol GP, Spreafico R, Wong V, Scott-Browne JP, Togher S, Hoffmann A, Hogan PG, Rao A, Trifari S. Exhaustion-associated regulatory regions in CD8(+) tumor-infiltrating T cells. Proc Natl Acad Sci U S A 2017;114(13):E2776–E85.

47. Foster SL, Hargreaves DC, Medzhitov R. Gene-specific control of inflammation by TLR-induced chromatin modifications. Nature 2007;447(7147):972–8.

48. Vire E, Brenner C, Deplus R, Blanchon L, Fraga M, Didelot C, Morey L, Van Eynde A, Bernard D, Vanderwinden JM, et al. The Polycomb group protein EZH2 directly controls DNA methylation. Nature 2006;439(7078):871–4.

49. Yaseen I, Kaur P, Nandicoori VK, Khosla S. Mycobacteria modulate host epigenetic machinery by Rv1988 methylation of a non-tail arginine of histone H3. Nat Commun 2015;6:8922.

50. Pennini ME, Pai RK, Schultz DC, Boom WH, Harding CV. Mycobacterium tuberculosis 19-kDa lipoprotein inhibits IFN-gamma-induced chromatin remodeling of MHC2TA by TLR2 and MAPK signaling. J Immunol 2006;176(7):4323–30.

51. Marr AK, MacIsaac JL, Jiang R, Airo AM, Kobor MS, McMaster WR. Leishmania donovani infection causes distinct epigenetic DNA methylation changes in host macrophages. PLoS Pathog 2014;10(10):e1004419.

52. Durzynska J, Lesniewicz K, Poreba E. Human papillomaviruses in epigenetic regulations. Mutat Res Rev Mutat Res 2017;772:36–50.

53. Chandran A, Antony C, Jose L, Mundayoor S, Natarajan K, Kumar RA. Mycobacterium tuberculosis Infection Induces HDAC1-Mediated Suppression of IL-12B Gene Expression in Macrophages. Front Cell Infect Microbiol 2015;5:90.

54. Mandalakas AM, Kirchner HL, Lombard C, Walzl G, Grewal HM, Gie RP, Hesseling AC. Well-quantified tuberculosis exposure is a reliable surrogate measure of tuberculosis infection. Int J Tuberc Lung Dis 2012;16(8):1033–9.

55. Liberzon A, Birger C, Thorvaldsdottir H, Ghandi M, Mesirov JP, Tamayo P. The Molecular Signatures Database (MSigDB) hallmark gene set collection. Cell Syst 2015;1(6):417–25.

56. Reinius LE, Acevedo N, Joerink M, Pershagen G, Dahlen SE, Greco D, Soderhall C, Scheynius A, Kere J. Differential DNA methylation in purified human blood cells: implications for cell lineage and studies on disease susceptibility. PLoS One 2012;7(7):e41361.

57. Lindqvist BM, Wingren S, Motlagh PB, Nilsson TK. Whole genome DNA methylation signature of HER2-positive breast cancer. Epigenetics 2014;9(8):1149–62.

